# Imaging-Guided Bioreactor for De-Epithelialization and Long-Term Cultivation of *Ex Vivo* Rat Trachea

**DOI:** 10.1101/2021.12.06.470803

**Authors:** Seyed Mohammad Mir, Jiawen Chen, Meghan R. Pinezich, John D. O’Neill, Sarah X.L. Huang, Gordana Vunjak-Novakovic, Jinho Kim

**Affiliations:** Department of Biomedical Engineering, Stevens Institute of Technology, Hoboken, NJ, USA; Department of Biomedical Engineering, Columbia University, New York, NY, USA; Department of Cell Biology, State University of New York Downstate Medical Center, Brooklyn, NY, USA; Center for Stem Cell and Regenerative Medicine, University of Texas Health Science Center, Houston, TX, USA

## Abstract

Recent synergistic advances in organ-on-chip and tissue engineering technologies offer opportunities to create *in vitro*-grown tissue or organ constructs that can faithfully recapitulate their *in vivo* counterparts. Such *in vitro* tissue or organ constructs can be utilized in multiple applications, including rapid drug screening, high-fidelity disease modeling, and precision medicine. Here, we report an imaging-guided bioreactor that allows *in situ* monitoring of the lumen of *ex vivo* airway tissues during controlled *in vitro* tissue manipulation and cultivation of isolated rat trachea. Using this platform, we demonstrated selective removal of the rat tracheal epithelium (i.e., de-epithelialization) without disrupting the underlying subepithelial cells and extracellular matrix. Through different tissue evaluation assays, such as immunofluorescent staining, DNA/protein quantification, and electron beam microscopy, we showed that the epithelium of the tracheal lumen can be effectively removed with negligible disruption in the underlying tissue layers, such as cartilage and blood vessel. Notably, using a custom-built micro-optical imaging device integrated with the bioreactor, the trachea lumen was visualized at the cellular level in real time, and removal of the endogenous epithelium and distribution of locally delivered exogenous cells were demonstrated *in situ*. Moreover, the de-epithelialized trachea supported on the bioreactor allowed attachment and growth of exogenous cells seeded topically on its denuded tissue surface. Collectively, the results suggest that our imaging-enabled rat trachea bioreactor and selective cell replacement method can facilitate creating of bioengineered *in vitro* airway tissue that can be used in different biomedical applications.

## Introduction

The human conducting airways are lined by the airway epithelium that mostly consists of multi-ciliated, club, goblet, and basal cells.^1–3^ These airway epithelial cells collectively create a protective biophysical barrier between the external environment and underlying tissues against inhaled harmful substances, such as pathogens, allergens, chemical gases, or particulates.^4^ The protective functions of the airway epithelium include mucociliary clearance,^5^ tight junction formation,^6^ and antimicrobial and anti-inflammatory secretions.^7^ In addition to serving as the first line of defense of the lung, the airway epithelium is the prime site for the initiation and progression of many devastating respiratory disorders. For example, cystic fibrosis (CF), primary dyskinesia (PCD), and chronic obstructive pulmonary disease (COPD) are diseases caused by genetic mutations or environmental insults that primarily impact the airway epithelium, contributing to functional decline of the lung and ultimate end-stage lung diseases.^8–10^ In CF patients, mutations in the cystic fibrosis transmembrane conductance regulator (CFTR) gene result in abnormal accumulation of viscous mucus within the airways and subsequent chronic airway infection and injury.^8,11^ Moreover, PCD is caused by mutations of proteins involved in cilia structure, organization, and function, and it is characterized by chronic infection of the upper and lower airways due to ineffective airways clearance by ciliated cells.^9,12^ Furthermore, the prolonged exposure of airway epithelium to noxious particles or gases can lead to COPD, accompanied by goblet cell hyperplasia, cell metaplasia, impaired ciliary function, mucus hypersecretion, and fibrotic tissue development.^10,13^

Recent synergistic advances in tissue engineering, biomaterials, and stem cell technologies have shown great potential for modeling different lung diseases,^14^ repairing damaged tissues or organs,^15^ studying human lung development,^16^ and developing therapeutic materials without the use of animals.^17^ For example, faithful *in vitro*models of diseases, such as cystic fibrosis,^18,19^ COPD,^20^ asthma,^21^ surfactant protein B deficiency,^22^ pulmonary fibrosis,^23,24^ and viral infection,^25^ have been generated using primary epithelial cells and induced pluripotent stem cells (iPSCs). Furthermore, recent establishment of innovative protocols for guided differentiation of airway stem cells, in particular basal cells,^26–28^ allowed emergence of cell replacement therapy as a promising approach to repairing airway tissues of live patients that are severely injured or diseased beyond their intrinsic repairable limits,^29^ or donor lungs that are refused for transplantation due to substantial tissue damage.^30^

To study differentiation and therapeutic functionality of airway stem cells, various *in vitro* platforms have been created and used. Traditionally, airway stem cells have been cultured and assessed in a static two-dimensional (2D) environment of a petri dish or transwell insert that can provide controllable cell culture conditions.^31–33^ The 2D-cultured cell monolayer enabled fundamental studies related to cell signaling pathways, cellular responses, and cell differentiation.^34,35^ Notably, lung-on-a-chip (LOC) devices with lung-mimetic designs have been developed that can allow co-culturing of different lung cells in an environment where fluids (e.g., air and culture media) are dynamically manipulated.^36^ For instance, airway epithelium and endothelium were cultured on the apical and basal sides of a porous membrane, respectively, to mimic breathing airway, allowing *in vitro* drug screening and disease modeling.^37–39^ However, these *in vitro* platforms are incapable of recapitulating the complex three-dimensional (3D) architecture and dynamic cell-matrix interactions found *in vivo*, resulting in considerable differences between 2D-cultured cells and *in vivo* cells in terms of morphology, proliferation, and differentiation.^40,41^

Decellularized allogeneic or xenogeneic tissue grafts have been used to provide *in vivo*-like microenvironments to the airway epithelial cells or stem cells during cell culture.^42,43^ For example, isolated rat or mouse tracheas with their endogenous cellular components completely removed via repeated freezing and thawing or chemical treatments were seeded with airway cells.^44,45^ When the cell-seeded decellularized airway tissues were implanted subcutaneously into immunodeficient host animals,^43^functional airway epithelial layer was regenerated on the luminal surface of the tissues. The *in vivo* cultured tissuespecific scaffolds allowed study of stem cell differentiation and confirmed regenerative capacity of the airway stem cells.^46^ Because cell-seeded tissue scaffolds are embedded in the host body, however, major limitations of this tissue culture approach include lack of ability for real-time manipulation of the cell culture conditions and monitoring of the cellular responses. In particular, creation of the air-liquid interface and time-dependent supply of growth factors or cytokines that are essential for stem cell differentiation are difficult to achieve in the *in vivo*-cultured tissue scaffolds.^47^ Further, different from *in vitro*-cultured models, microscopic assessments of the cultured cells are only possible after removing the cell-tissue constructs from the host after completion of each experiment.^44,48^

Here, we report an imaging-enabled rat lung bioreactor system that allows long-term *in vitro* cultivation of isolated rat trachea and direct visualization of the tracheal lumen at the cellular level (**Fig. 1**). Using this platform, we demonstrated selective removal of the epithelial layer from the trachea lumen without disrupting the underlying tissue layers and extracellular matrix (ECM) components, onto which exogeneous airway cells or stem cells could beimplanted to restore functional epithelium. Further, we created a micro-imaging device that can be inserted into the inner space of the *in vitro*-cultured trachea to allow *in situ* visual monitoring of the cell replacement and cultivation. Notably, the de-epithelialized rat tracheas supported survival of exogenous cells, such as mesenchymal stem cells (MSCs), that were topically seeded onto the denuded tracheal lumen. We envision that the imaging-enabled bioreactor platform and tissue manipulation methodology established in this study could enable creation of humanized airway tissues by combining with human airway stem cells, allowing *in vitro* study of airway diseases and expediting development of therapeutics and intervention modalities.

**Fig. 1.**
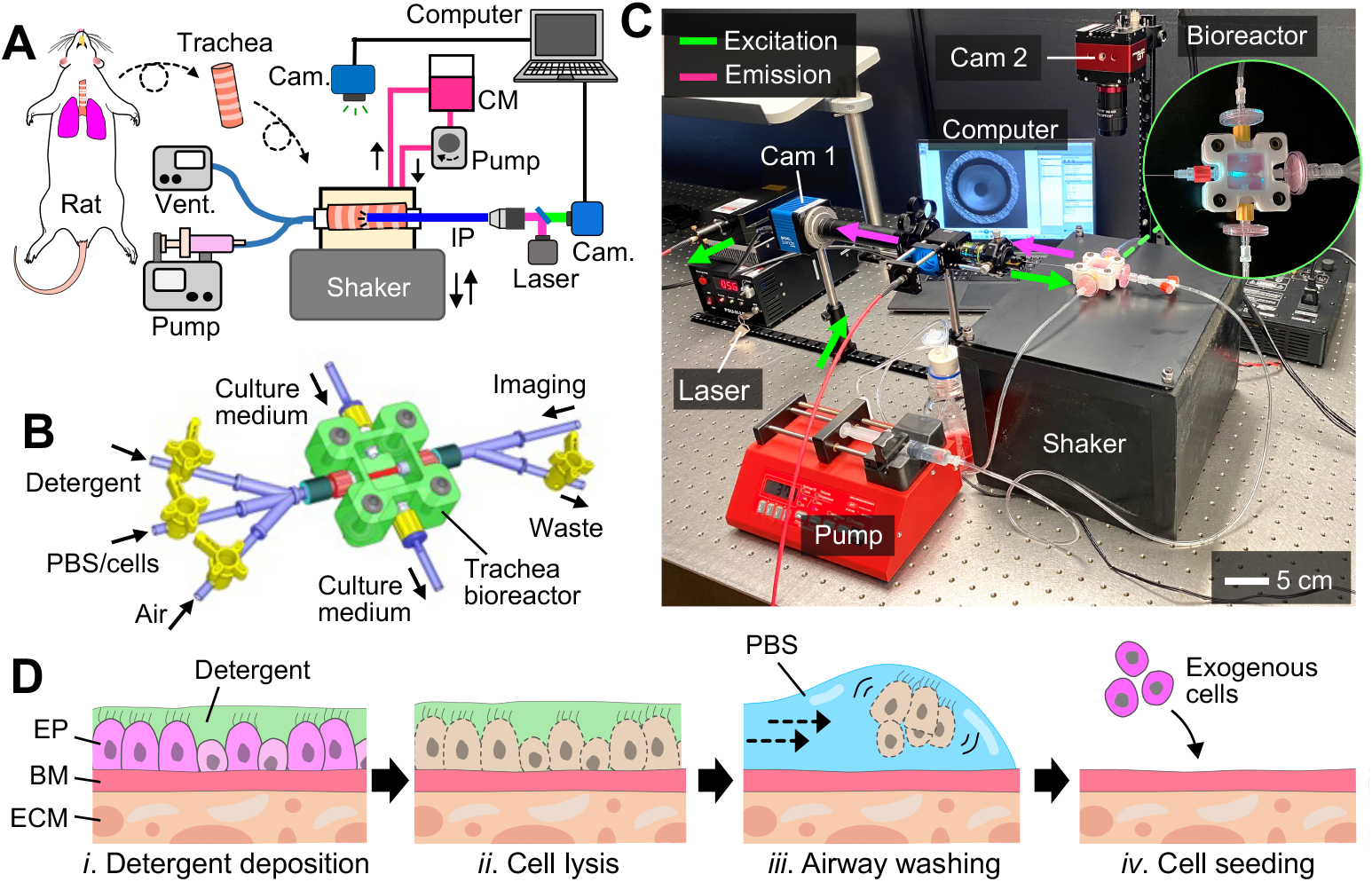
Overview of imaging-enabled bioreactor and de-epithelialization of *ex vivo* rat trachea: (**A**) Schematic of the imaging-integrated rat trachea bioreactor platform. Cam: camera. CM: culture medium. IP: imaging probe. Vent: ventilation. (**B**) Three-dimensional (3D) representation of the trachea bioreactor. (**C**) Photograph of the bioreactor system. Cam: camera. (**D**) Schematic showing the procedure of de-epithelialization of rat trachea. EP: epithelium. BM: basement membrane. ECM: extracellular matrix. PBS: phosphate-buffered saline.

## Results

### Imaging-enabled bioreactor for *in vitro* cultivation of isolated rat trachea

We constructed an imaging-enabled bioreactor system that enabled selective removal of the epithelium and long-term culture of the tracheal tissue (**Fig. 1A**). The bioreactor was designed and constructed in a way that the luminal surface of the trachea can be treated using different solutions (e.g., decellularization solution, washing solution, cells, culture medium) while the entire trachea is submerged in a cell culture medium to maintain the viability of the trachea tissue during planned experiments (**Fig. 1B, C**). Using this system, isolated rat trachea can be de-epithelialized by introducing decellularization reagents (i.e., sodium dodecyl sulfate detergent; SDS) with specified volumes and concentrations directly into the inner space of the trachea. Disrupted cells can be removed from the tissue surface with phosphate-buffered saline (PBS) washing solution. The airway imaging device integrated into the bioreactor enables direct *in situ* visual inspection of the trachea lumen in both bright-field and fluorescence modes. While it was not demonstrated in this current study, the trachea could be supplied with a gas flow (e.g., air) via the ventilation port connected to the bioreactor to create air-liquid interface within the trachea or to provide *in vivo*-like flow shear stress to the seeded cells.

### De-epithelialization of *ex vivo* rat trachea

To selectively remove the tracheal epithelium without disrupting the rest of the underlying airway tissue (**Fig. 1D**), we deposited a thin layer of 2% or 4% SDS detergent topically onto the tracheal lumen by instilling a small volume (50 μL) of the detergent solution. Instillation of a small volume of liquid (e.g., aqueous solution) through the respiratory tract can generate a thin layer of the liquid onto the airway lumen.^49–51^ Following disruption of the epithelium through detergent exposure, the lysed cells were then cleared from the trachea by washing with PBS buffer while the entire trachea was vibrated mechanically using our custom-built shaker (**Fig. S1**). In particular, the trachea was oscillated at 20 Hz with a vertical displacement of approximately 0.3 mm (**Video S1** and **Video S2**). Exogeneous cells (e.g., airway epithelial or stem cells) can be seeded onto the denuded tracheal lumen to reconstitute the epithelium within the bioreactor.

### *In situ* visualization of the trachea lumen

We used our custom-built in situ airway imaging device to inspect the luminal surface of the rat trachea tissue during *in vitro* cell removal (**Fig. 2A, *i-iii***). We used a 1951 USAF (U.S. Air Force) test target to evaluate the resolution of the images obtained using the device. Obtained images showed high resolution and good quality with approximately 5 μm of the smallest features resolvable (**Fig. 2A, *iv***). To visualize the trachea lumen, the imaging probe was directly inserted into the trachea through a cannula connected to the trachea (**Fig. 2B, *i-ii***). Both bright-field (**Fig. 2C, *i*; Video S3**) and fluorescence images (**Fig. 2C, *ii***) of the local luminal surface were obtained, respectively, by using white light and 488-nm laser for illumination prior to carboxyfluorescein succinimidyl ester (CFSE) labeling of the epithelial layer. While no fluorescence signal was observed before labeling, a discernible signal (i.e., green light) was observed when the epithelium was labeled with CFSE (**Fig. 2D, *i*** and **Video S4**). Notably, following de-epithelialization, the intensity of the fluorescent signal substantially decreased, indicating clearance of the epithelium from the tracheal lumen (**Fig. 2D**, ***ii***). This result highlights the utility of our imaging approach in *in situ* and minimally invasive monitoring of the epithelium removal.

**Fig. 2.**
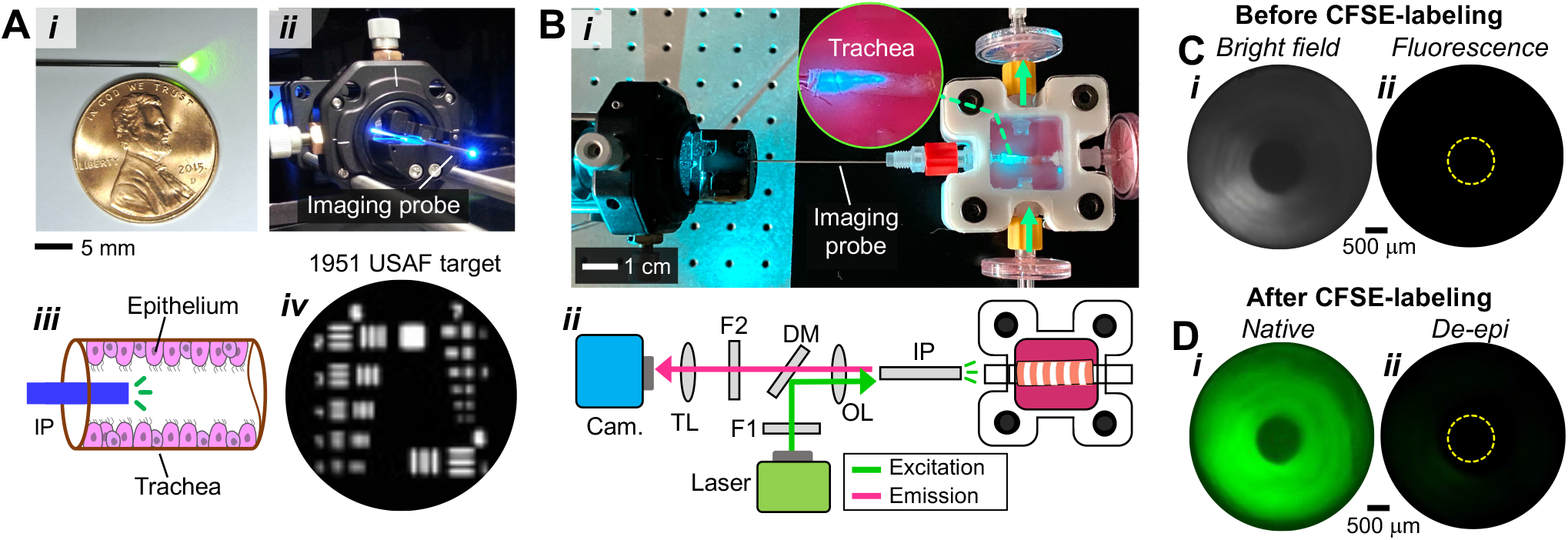
*In situ* visualization of the trachea lumen using custom-built micro-optical imaging device: (**A**) (*i-iii*) GRIN lens (diameter: 500 μm) used for both bright-field and fluorescence imaging of the rat tracheal lumen. (*iv*) 1951 USAF test target imaged using the imaging device. (**B**) (*i*) Photograph and (*ii*) schematic showing the imaging probe being used for visual inspection of an *in vitro*-cultured rat trachea. Cam: camera. TL: tube lens. F: optical filter. DM: dichroic mirror. OL: objective lens. IP: imaging probe. (**C**) (*i*) Bright-field and (*ii*) fluorescence images of the interior of the rat trachea before CFSE-labelling of the epithelium. (**D**) Fluorescence images of (*i*) native and (*ii*) de-epithelialized (De-epi) rat trachea lumen that was labelled with CFSE.

### Selective removal of tracheal epithelium with preserved tissue ECM

Histological evaluation of de-epithelialized tracheas showed complete removal of the epithelium across the luminal surface of de-epithelialized tracheas treated with 2% and 4% detergent solutions (**Fig. 3**). In particular, high magnification of H&E images confirmed the removal of the pseudostratified columnar epithelium from the trachea mucosa and preservation of other cells in the submucosa, cartilaginous, and adventitia layers (**Fig. 3A** and **Fig. S2**). In addition, the structure pattern of the tissue layers and endogenous cells, such as cartilage and chondrocytes, underneath the basement membrane was well preserved following de-epithelialization. Similarly, pentachrome (**Fig. 3B**) and trichrome (**Fig. 3C**) staining revealed maintenance of the tissue architecture and ECM components, including collagen and proteoglycans (e.g., mucins), while the airway epithelium was cleared from the lumen.

**Fig. 3.**
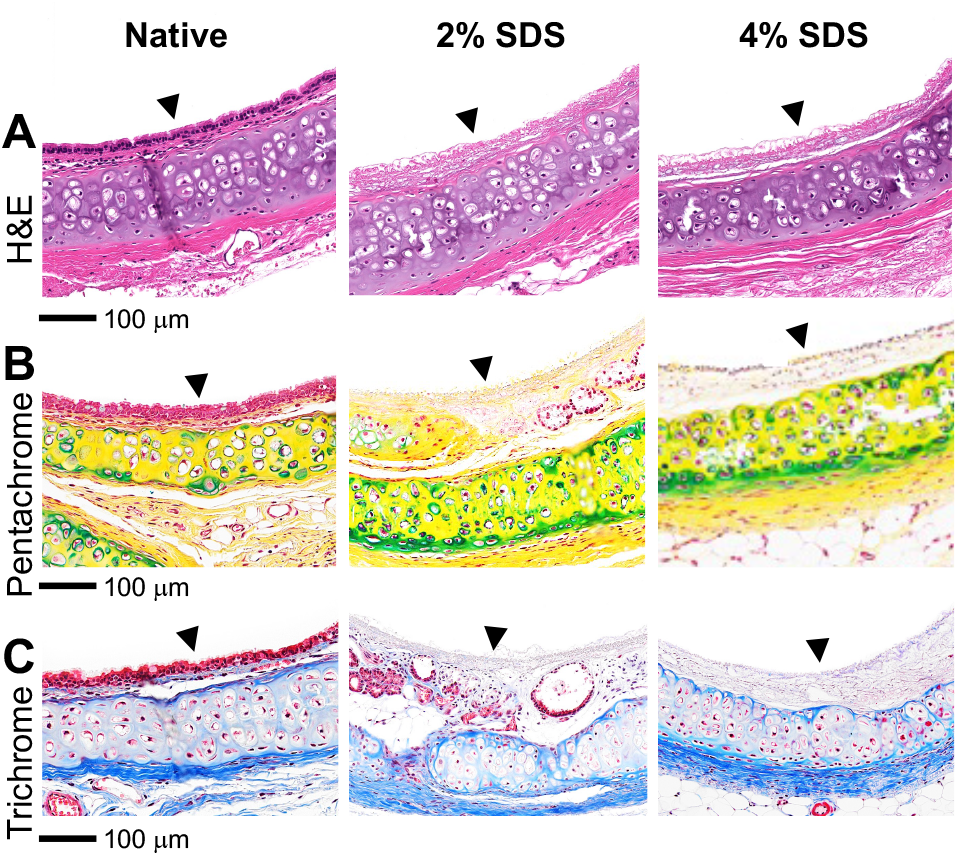
Histologic analysis of native and de-epithelialized rat tracheas: (**A**) H&E, (**B**) pentachrome, and (**C**) trichrome staining images of native and de-epithelialized rat tracheas with 2% and 4% SDS. The images of de-epithelialized tracheas show removal of the epithelium and preservation of ECM architecture. *Pentachrome: purple (cell cytoplasm), blue (cell nuclei), green (proteoglycans), yellow (collagen fibers). *Trichrome: pink (cell cytoplasm), dark blue (cell nuclei), blue (collagen). Arrowhead: tracheal lumen.

Further, we confirmed the preservation of ECM components of the de-epithelialized tracheas via immunofluorescence staining. Immunostaining of the trachea tissues by epithelial cell adhesion molecule (EpCAM) and 4’,6-diamidino-2-phenylindole (DAPI) revealed removal of the epithelial layer as no EpCAM (green) and DAPI (blue) signals were detected at the lumen of the trachea (**Fig. 4A**). De-epithelialized tracheas treated with 4% SDS showed a reduction in the DAPI signal throughout the tissue, suggesting potential disruptions occurred to endogenous subepithelial cells due to increased detergent concentration. Nevertheless, the de-epithelialization method preserved laminin (green) (**Fig. 4B**), collagen I, elastin, and smooth muscles of the trachea tissue (**Fig. S3**). Furthermore, immunostaining of the de-epithelialized tissues with endothelial cell marker (cluster of differentiation 31; CD31) confirmed that the blood vessels of the tissue remained intact (**Fig. 4C**).

**Fig. 4.**
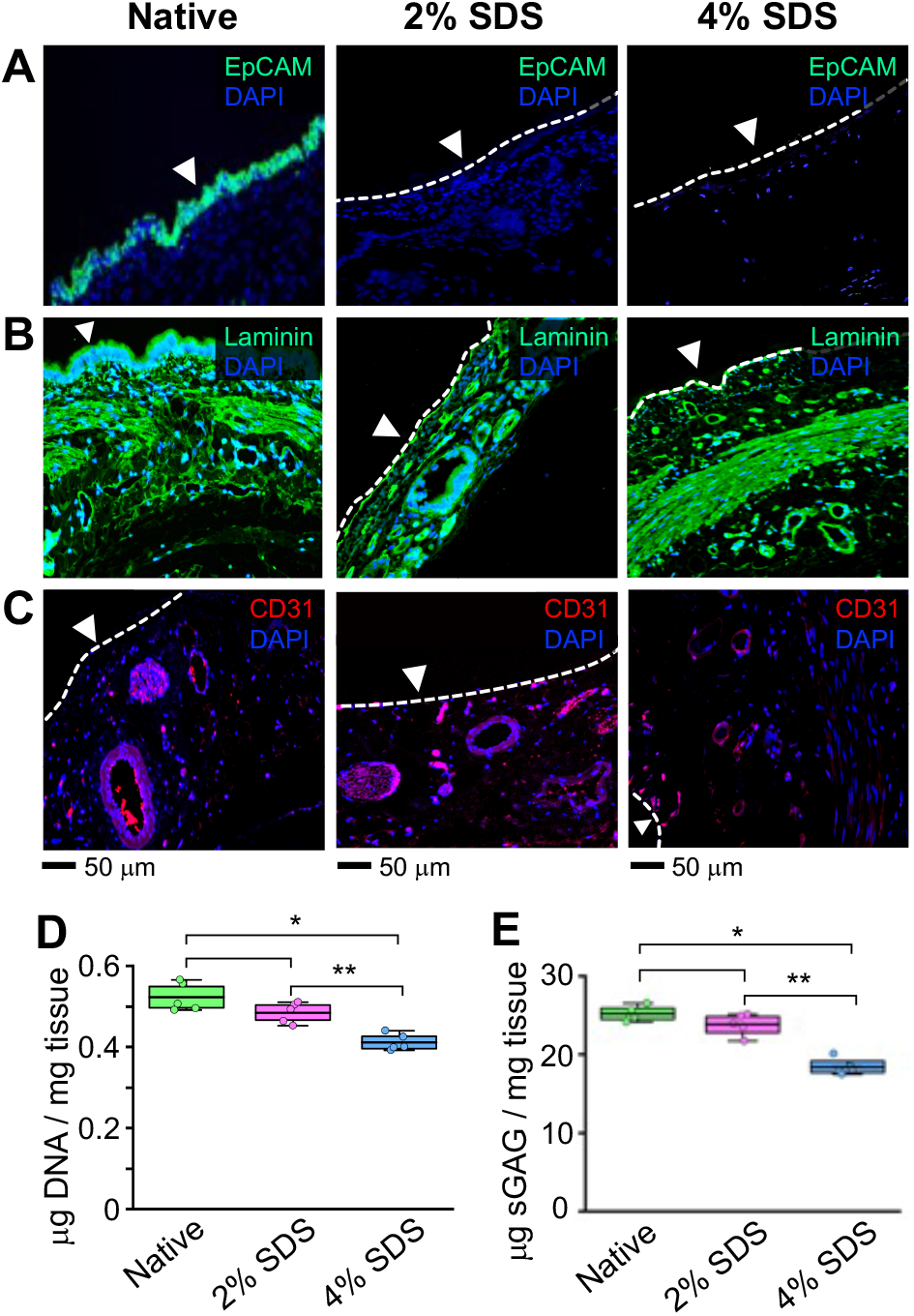
Evaluation of ECM components via immunohistochemistry and DNA/GAG quantification: Immunostaining images of (**A**) epithelial cell adhesion molecule (EpCAM), (**B**) laminin, and (**C**) cluster differentiation 31 (CD31) showing removal of the tracheal epithelium and preservation of ECM components within the rat trachea treated with 2% and 4% SDS. Arrowhead: tracheal lumen. (**D**) DNA and (**E**) GAG quantifications (*n* = 5 each) decrease in DNA and sulfated GAG amounts. Error bars represent means ± SD of experimental values. \**p* < 0.001. \*\**p* < 0.01.

DNA quantification showed no significant differences between native tissues (0.52 ±0.03 μg.mg^-1^ of tissue) and de-epithelialized tracheas treated with 2% SDS (0.48 ±0.03 μg.mg^-1^ of tissue; *p* = 0.11) (**Fig. 4D**). On the other hand, the DNA content decreased by nearly 21% in the tracheas treated with 4% SDS (0.41 ±0.02 μg.mg^-1^ of tissue; *p*< 0.001), indicating some of the endogenous cells were affected when the trachea tissues were exposed to the higher concentration of the detergent solution. Furthermore, the glycosaminoglycan (GAG) quantification assay (**Fig. 4E**) revealed no significant difference in sulfated GAG content between native trachea (25.2 ±0.9 μg.mg^-1^ tissue) and de-epithelialized tracheas treated with 2% SDS solution (23.8 ±1.4 μg.mg^-1^ of tissue; *p*= 0.17). However, similar to DAPI staining and DNA quantification, nearly 27% decrease in the sulfated GAGs was observed in the tracheas treated with 4% SDS solution (18.4 ±1.0 μg.mg^-1^ of tissue; *p* < 0.001).

### Scanning electron microscopy (SEM) imaging of the tracheal lumen

We then investigated topological changes in the luminal surfaces of de-epithelialized trachea via SEM imaging where the images were obtained at different magnifications (30×in **Fig. 5A**; 2000×in **Fig. 5B**; and 4000×in **Fig. 5C**). The native trachea showed that the luminal surface was densely populated by different epithelial cells, predominantly multi-ciliated and goblet cells. On the other hand, the SEM images of de-epithelialized tracheas with 2% and 4% SDS showed absence of the tracheal epithelium. Notably, in both de-epithelialized trachea lumen surfaces, a thin membrane layer, which is most likely the basement membrane, and mesh network of airway ECM were clearly visible. Structural disruption of the basement membrane and tissue ECM was more prominent in the tracheas treated with 4% SDS compared with that of 2%SDS as the porosity of the remaining ECM structure increased with the concentration of the SDS. Notably, use of mechanical vibration during the airway washing facilitated detachment of the epithelium as SDS-disrupted epithelium remained attached onto the lumen surface when no vibration was applied to the tissue (**Fig. 4S**). This result clearly indicated that oscillation energy provided to the trachea in the presence of the shear flow promoted disruption and detachment of the detergent-lysed cells from the airway tissue ECM.

**Fig. 5.**
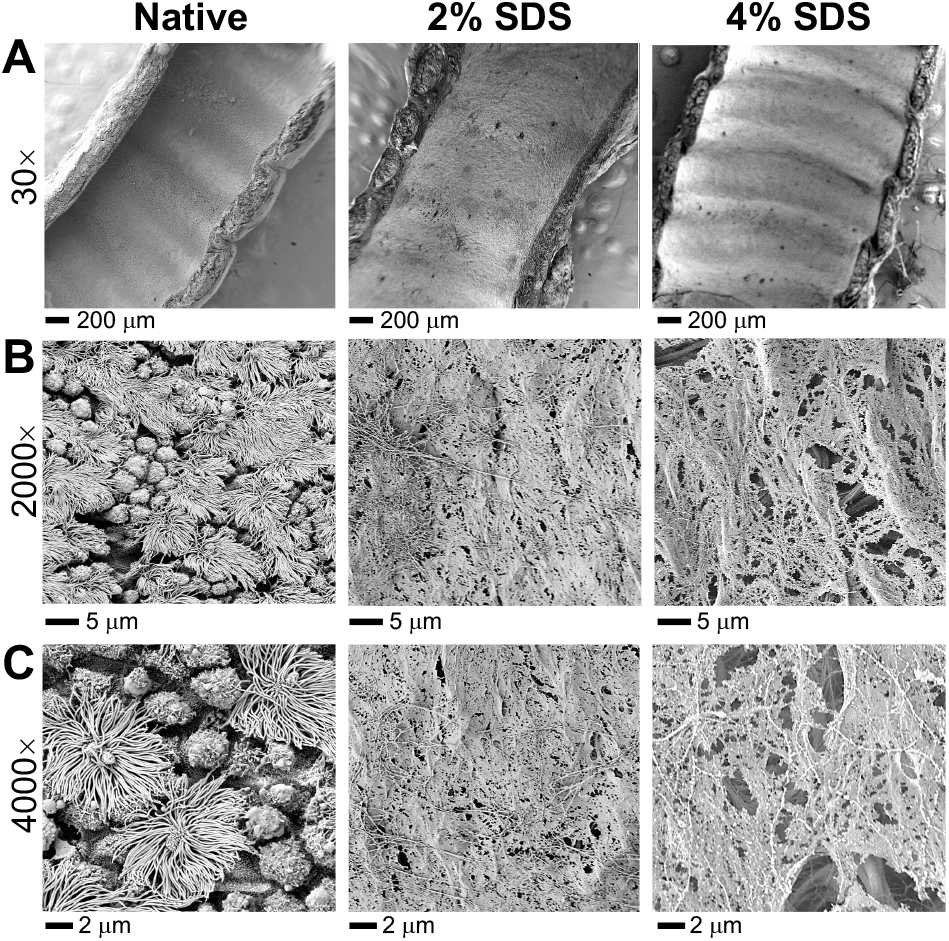
Investigation of the luminal surface of de-epithelialized tracheas via SEM imaging: SEM images obtained at (**A**) 30×, (**B**) 2ŪÜÜ×, and (**C**) 4ÜÜÜ×of magnifications showing the luminal surface of the *ex vivo* rat tracheas.

### Evaluation of chondrocyte viability and cytotoxicity of the de-epithelialized trachea tissue

We assessed the viability of chondrocytes in native and de-epithelialized tracheas after culturing for two weeks in the bioreactor. We used the de-epithelialized trachea treated with 4% SDS solution, which is the maximum concentration of SDS used in this study. The fluorescent images of the trachea sections revealed that most of the chondrocytes seeded onto native and de-epithelialized tracheas survived following two weeks of the *in vitro* cultivation (**Fig. 6A**). In addition, quantification of viable cartilage cells (green) and the dead cells (red) showed that both native (86.2%±1.4%) and de-epithelialized (84%± 1.8%) tracheas maintained the viability of majority of the cartilage cells (**Fig. 6B**).

**Fig. 6.**
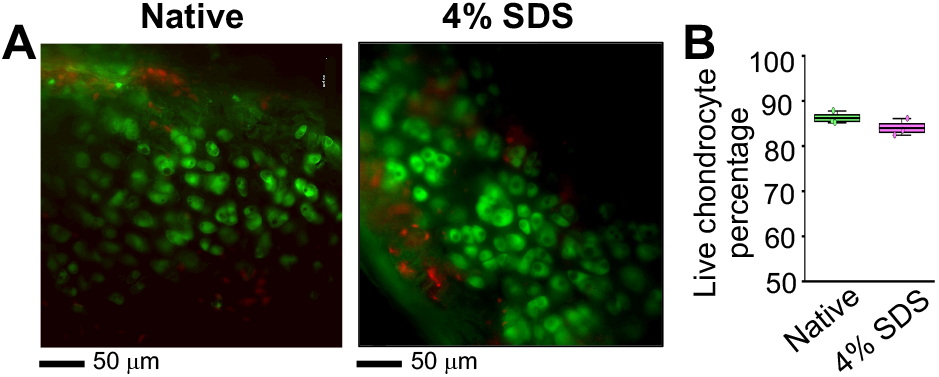
Live and dead analysis of native and de-epithelialized trachea (4% SDS): (**A**) fluorescent images of live and dead cartilage cells showing the viability of chondrocytes after de-epithelialization and two weeks of *in vitro* cultivation. (**B**) Quantification of the viable chondrocyte revealed that there was no significant difference between native and de-epithelialized tracheas (*p* = 0.18).

We also investigated whether the SDS-treated tracheas can support attachment, growth, and proliferation of exogenous cells seeded on the lumen surface without generating a cytotoxic environment. To do this, CFSE-labelled MSCs were introduced and topically deposited onto the inner space of *ex vivo* rat tracheas with their epithelium removed via 4% SDS (**Fig. 7**). Immediately after cell seeding, the micro-imaging probe was inserted into the trachea to confirm the deposition of the cells (green) onto the de-epithelialized tracheal lumen (**Fig. 7A, *i-ii***). The bioreactor containing the cell-seeded trachea was then transferred to an incubator and connected to the perfusion pumps for extended *in vitro* culture (i.e., 1, 4, and 7 days) (**Fig. 7B, C**). The fluorescent images of the MSCs cultured on the de-epithelialized tracheal lumen showed that the density of the cells covering the surface gradually increased over time (i.e., day 1: 15.6 ±6.1 cells.mm^-2^, day 4: 94.6 ±15.1 cells.mm^-2^, day 7: 145 ±12.1 cells.mm^-2^; **Fig. 7D, E**). Further, the average circularity of the seeded cells, which is a normalized ratio of area to perimeter of the cells, was calculated to quantitatively evaluate attachment and engraftment of the cells on the surface. Analysis showed that cell circularity decreased over time, confirming active binding and incorporation of the cells onto the de-epithelialized tracheal lumen (i.e., day 1: 0.90 ±0.07, day 4: 0.22 ±0.04, day 7: 0.25 ±0.08; **Fig. S5**). Notably, SEM imaging showed that seeded MSCs quickly initiated binding onto de-epithelialized lumen at day 1 as multiple protrusions extending from the cell membrane to the tissue surface were observed (**Fig. S6**). Collectively, the results indicated that the de-epithelialized trachea lumen provided a suitable microenvironment that allowed engraftment, survival, and proliferation of topically seeded exogenous cells.

**Fig. 7.**
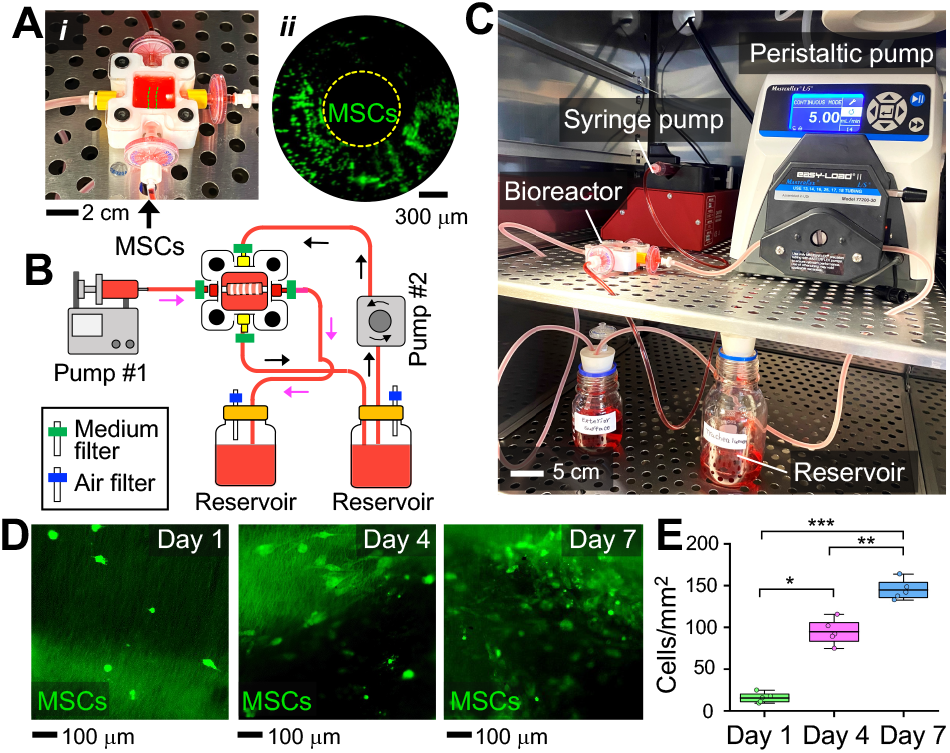
Evaluation of cytotoxicity of the rat tracheas de-epithelialized via 4% SDS detergent: (**A**) (*i*, *ii*) CFSE-labelled MSCs (green) were deposited onto the denuded rat trachea lumen using the micro-optical imaging device. (**B**) Schematic and (**C**) photograph showing the experimental setup used for long-term *in vitro* culture (i.e., over one week) of MSC-seeded de-epithelialized rat tracheas. Arrows show flow directions. (**D**) Fluorescence images of CFSE-labelled MSCs (green) being cultured on the tissue surface over the course of 7 days. (**E**) Density of the cells measured at different time points. Error bars represent means ± SD of experimental values. \**p* < 0.01. \*\**p* < 0.001. \*\*\**p* < 0.0001

## Discussion

To promote survival, proliferation, and differentiation of implanted exogenous cells, preservation of the underlying ECM structure and components within decellularized tissue is critical. The ECM provides biochemical and mechanical cues as well as suitable physical environments that are essential for tissue- or organ-specific cellular responses and activities.^52–55^ Recent studies demonstrated that airway basal stem cells implanted into decellularized trachea tissues are able to reconstitute a wide range of airway epithelial cell types, including multi-ciliated and secretory cells, that were difficult to achieve using traditional 2D cell culture platforms.^26,28,42,47^ On the other hand, the limitations of existing methods of culturing stem cell-implanted airway scaffolds include lack of methods to precisely modulate cell culture conditions and rapidly evaluate the behaviour and responses of implanted cells. Because cell-seeded tissue scaffolds are typically implanted in the host animal’s body during the culture, controlling and monitoring the cells and tissues *in situ* are very challenging.^56,57^ Also, existing decellularization protocols focus on the complete removal of all existing cellular components from the airway tissues (e.g., epithelium, endothelium, stromal cells, etc.), which eliminates the possibility of investigating interactions between resident subepithelial cells and newly implanted stem cells.^58–60^

In this work, we developed an imaging-enabled bioreactor platform that allows long-term culture (at least one week) of cell-implanted rat trachea in a precisely modulated *in vitro* environment (**Fig. 1**). This platform is capable of: (*i*) applying user-defined biochemical treatments to *ex vivo* airway tissue; (*ii*) *in situ* monitoring of the interior and exterior surfaces of the airway at the cellular level; and (*iii*) long-term culturing of *ex vivo* tissues following topical seeding of exogenous cells. Further, we developed a tissue-manipulation protocol that permitted selective removal of the epithelium of the trachea lumen while preserving the rest of the tissue layers and ECM components. This targeted epithelium removal approach utilizes topical deposition of a liquid solution containing decellularization reagents (e.g., detergent solution) directly onto the airway lumen. By modulating the decellularization reaction time and detergent strength, the epithelium can be dislodged and removed with washing solution under mechanical vibration. Vibration-assisted airway washing used in our study can further reduce the detergent amount and reaction time needed to lyse the epithelium, minimizing detergent-induced damage and detergent residue within the ECM.

Using this innovative platform and protocol, we demonstrated imaging-guided removal of the epithelial layer from the lumen of isolated rat trachea where fluorescently labeled epithelium was effectively ablated from the tissue surface. Notably, the cell removal was monitored visually *in situ* using our custom-built imaging device incorporated into the bioreactor (**Fig. 2**). We further confirmed the selective epithelium removal through additional tissue assessment methods, including histology, immunostaining, DNA/protein quantification, and electron beam microscopy (**Fig. 3**, **Fig. 4**, and **Fig. 5**). All tissue evaluation results consistently indicated that the tracheal epithelium could be selectively ablated while the ECM components and the rest of the cells underneath the basement membrane of the treated trachea could be preserved using our targeted epithelium removal approach. Higher concentration of detergent (4% SDS vs. 2% SDS) appeared to increase disruption in the ECM slightly. The rat tracheas treated with 4% SDS, compared with the tracheas treated with 2% SDS, resulted in the decreased number of subepithelial cells (**Fig. 4A**), reduced amounts of DNA and GAGs (**Fig. 4B**), while ECM structure became more porous (**Fig. 5**). Nevertheless, long-term cell culture experiments showed that rat tracheas de-epithelialized using 4% SDS detergent preserved most of the cartilage cells and were able to support active engraftment and proliferation of exogenous MSCs seeded onto the ECM surface (**Fig. 6** and **Fig. 7**).

The experimental results suggested that our imaging-enabled rat trachea bioreactor can allow the creation of bioengineered trachea tissue constructs. For example, in the future, human airway basal stem cells can be implanted onto de-epithelialized rat tracheas to reconstitute functional epithelium using our platform to generate *in vitro* humanized airway tissues. Such humanized airway constructs could potentially be used to investigate human airway diseases, screen therapeutics, and study human stem cell differentiation without the need of costly immunodeficient animals that are difficult to maintain and manipulate. In addition, future studies should evaluate the creation of the air-liquid interface (ALI) within the cultured trachea in our bioreactor to guide the differentiation of implanted basal cells into different epithelial cell types. The creation of ALI can be achieved by selectively controlling the infusion of culture medium and airflow through dedicated inlet ports.

Further, the gentle de-epithelialization methodology can be modified and potentially translated into clinics for “stem cell implantation therapy.” For example, diseased or severely injured airway epithelium could be cleared from the airway tissue and patient-derived basal stem cells can be topically seeded to restore functional airway tissue. To translate this method, more rigorous studies will be required to determine the optimal concentration, amount, and incubation time of the de-epithelialization reagents (e.g., detergents, enzymes, etc.) within the geometrically complex human respiratory tract.^50^ The ability to locally deliver and monitor biochemical reagents or cells within the lung will be imperative to maximize the outcome of cell replacement therapy. Moreover, the *in situ* micro-imaging modality developed in this study could serve as a translational tool to evaluate newly delivered cells following therapeutic cell replacement in patients with airway disease.

## Conclusions

*In vitro* airway tissue constructs created in the laboratory can facilitate fundamental studies of stem cell differentiation, development of effective therapeutic materials, and investigation of initiation and resolution of respiratory disorders. Here, we presented an imaging-assisted bioreactor platform and tissue manipulation methodology that allow selective removal of the endogenous tracheal epithelium, topical implantation of exogenous cells, and subsequent *in vitro* culture of the *ex vivo* tissues for an extended period. In addition, the platform and modalities created in this study can contribute to expanding our knowledge of the dynamic interactions between implanted exogenous cells and native airway ECM by allowing precise modulation of culture conditions and *in situ* assessment of the cell-tissue constructs. Further, the methodologies developed can be potentially translated into the clinics following additional validations and refinements to accelerate establishment of stem cell-based treatments for lung diseases caused by dysfunctional airway epithelium.

## Methods

### Isolation and preparation of rat tracheas

In this study, tracheas isolated from Sprague-Dawley rats were used (total: 24 rats; weight: 250-270 g; SAS SD Rats, Charles River Laboratories). All procedures were performed in accordance with the animal welfare guideline and regulations of the Institute for Animal Care and Use Committee (IACUC) at Stevens Institute of Technology. Prior to lung isolation, the rats were anesthetized via inhalation of 2.5% isoflurane for 20 minutes using a vaporizer (SomnoSuite^®^, Kent Scientific). To prevent blood clotting in the pulmonary vasculature, 1,000 unit/mL heparin (1 mL) was injected through the lateral tail vein and euthanized the animal using an overdose of 5% isoflurane. Immediately after euthanasia, the trachea was cannulated using a tracheal cannula (diameter: 2 mm; Cat. No. 73-2727, Harvard Apparatus). Midline thoracotomy was performed to isolate the trachea from the animal. The trachea was washed with 20mL of 1× PBS at room temperature and was transferred into our custom-built bioreactor supplemented with culture medium containing Dulbecco’s modified Eagle’s medium (DMEM; Gibco^™^, Cat. No. 11965118, Thermo Fisher Scientific), recombinant human FGF-basic (Cat. No. 100-18B, PeproTech), fetal bovine serum (FBS; Gibco^™^, Cat. No. 10082147, Thermo Fisher Scientific), and antibiotic-antimycotic (Cat. No. 15240062, Thermo Fisher Scientific). The culture medium was used for both *in vitro* culture of cells and trachea tissues in this study.

### Construction of the trachea bioreactor

We created a rat trachea bioreactor for de-epithelialization and *in vitro* culture of the rat trachea. The bioreactor consisted of a trachea culture chamber (dimension: 2.5 cm ×2.5 cm × 1.5 cm; total volume: 9.4 mL) in which the isolated trachea was placed for de-epithelialization and subsequent *ex vivo* culture. The culture chamber was fabricated by machining autoclavable Teflon^®^ PTFE plastic (McMaster-Carr) using a computer numerical control (CNC) machine (Mini-Mill 3, Minitech^®^). Within the chamber, each end of the trachea was connected to a Luer cannula (diameter: 1.5 mm; Cat. No. EW-45501-30, Cole-Parmer). A multi-way connector was attached to the inlet cannula to apply different fluids and reagents to the trachea, such as detergent, washing solution, and culture medium. Through the outlet cannula, a Y-connector was attached to collect waste solution and introduce a micro-optical imaging probe for visualizing the inner surface of the trachea. A transparent acrylic plastic sheet (McMaster-Carr) was cut and attached to the top of the main chamber using button head screws (10-32 threads, diameter 4.76 mm, length: 3.175 cm; McMaster-Carr).

### Tracheal epithelium labeling

To enhance the visibility of the tracheal epithelium, the inner lumen of the trachea was labeled fluorescently using carboxyfluorescein succinimidyl ester (CFSE) cell labeling kit (ab113853, Abcam). The tissue labeling was performed according to the company’s instruction prior to the de-epithelialization of the trachea. Briefly, CFSE solution was diluted with 1× PBS and injected into the inner lumen of the trachea. The trachea was incubated for 10 min at 37°C and then washed with 1× PBS (volume: 10 mL) three times to remove non-incorporated CFSE from the luminal surface. Our custom-built imaging device was used for in situ monitoring of the CFSE-labelled epithelial cells (excitation/emission: 492 nm/518 nm).

### Custom-built micro-optical imaging device

To visualize the inner lumen of the trachea either in bright field or fluorescence, we built a micro-optical imaging device that could be integrated with the bioreactor. The imaging device comprised of a GRIN lens (diameter: 500 μm; SELFOC^®^, NSG Group), a scientific camera (PCO Panda 4.2), an achromatic doublet (AC254-150-A-ML, Thorlabs), a filter lens (ET535/50m, Chroma^®^), a dual-edge super-resolution dichroic mirror (DI03-R488, Semrock), and a 20× objective lens (UCPLFLN20×, Olympus). For imaging the tracheal lumen, the distal end of the GRIN lens was inserted into the Luer connector that was attached to one end of the trachea. The light signals emitted from the luminal surface of the trachea were detected at the proximal end of the lens, which were then visualized via the camera. For fluorescent imaging, a laser light (wavelength: 488 nm, power: 150 mW; MDL-D-488-150mW, Opto Engine) was passed into the imaging probe to excite the CFSE-labelled epithelium. Fluorescent light signals generated from the CFSE (emission wavelength: 515 nm) were filtered by the dichroic mirror and bandpass filter lens, then collected and imaged via the camera. The resolution of the imaging device was evaluated using a test target (1951 United States Air Force Resolution Test Target, Cat. No. R1DS1P, Thorlabs).

### Construction of an electromagnetic shaker

To facilitate the detachment of epithelial cells from the tracheal lumen, we constructed an electromagnetic shaker, onto which a trachea-loaded bioreactor can be placed and mechanically oscillated at a specified frequency and amplitude. The custom-built shaker was composed of a subwoofer speaker (RSS21OHO-4, Dayton Audio) and a subwoofer plate amplifier (SPA250DSP, Dayton Audio). A rigid acrylic plate (dimensions: 25 cm ×30 cm ×1.5 cm; McMaster-Carr) was attached onto the diaphragm of the speaker (diameter: 21 cm). The trachea bioreactor was loaded on the plate during mechanical oscillation. The frequency and amplitude of the shaker were manipulated by feeding a computer-generated sinusoidal waveform to the shaker. The response of the shaker was measured using an accelerometer (IIS3DWBTR, STMicroelectronics) or a high-frame-rate camera (240 frames per second; iPhone 11 Pro).

### De-epithelialization of *in vitro*-cultured rat trachea

To remove the epithelium from the *in vitro*-cultured rat trachea, an aqueous solution of sodium dodecyl sulfate (SDS; concentrations: 2% or 4%; Cat. No. 97064-472, VWR) was instilled directly into the trachea via an inlet cannula. Specifically, a small volume (50 μL) of either 2% or 4% SDS solution was infused through the trachea using a programmable syringe pump (flow rate: 6.3 mL.s^-1^; AL-4000, World Precision Instruments) to generate a thin film of the detergent solution on the luminal surface of the trachea.^49,51^ To promote de-epithelization, the trachea was incubated in the bioreactor for 20 min at 37°C. The bioreactor was then mechanically vibrated using our custom-built shaker at 20 Hz of frequency while being washed with 1× PBS solution three times (volume: 500 μL; flow rate: 10 mL.s^-1^).

### Histological analysis

All tracheas, native and de-epithelialized, were fixed in 4% neutral buffered paraformaldehyde solution in 1× PBS (pH level: 7.4) at 4°C overnight and transferred to a 70% ethanol solution. The fixed tracheas were then dehydrated in a series of isopropyl alcohol solutions (IPA; concentrations: 80, 90, 95, and 100%) and Citrisolv^™^ solution (Cat. No. 1061, Decon) prior to paraffin embedding. The paraffin-embedded trachea tissues were cut into thin slices (thickness: 5-8 μm) and were deparaffinized using Citrisolv^™^ solution followed by rehydration in a series of IPA solutions (concentrations: 100, 95, 90, 80, 70, 50%). For microscopic assessment of the tissue layers, the tissue sections were stained for hematoxylin and eosin (H&E), Masson’s trichrome, and Movat’s pentachrome at the Department of Molecular Pathology at Columbia University.

### Immunofluorescence analysis

The paraffin-embedded trachea sections were deparaffinized with Citrisolv^™^ and rehydrated in serial concentrations of 100 to 50% IPA solutions. To retrieve the antigens, the tissue section samples were treated with boiling citrate buffer at pH 6. The samples were then permeabilized in 1× PBS with 0.25% TritonTM X-100 (Cat. No. 97063-864, VWR) and FBS for 20 min. The tissue sections were then blocked in 10% FBS for additional 2 hours at room temperature and stained by incubating primary antibodies (**Table S1**) in a staining buffer containing 5% FBS and 1× PBS at 4°C overnight. The tissue sections were then washed three times in washing buffer and stained with corresponding secondary antibodies (**Table S2**). Lastly, the sections were washed, incubated with DAPI solution (4’,6-diamidino-2-phenylindole; ab228549, Abcam) for 15min, rewashed, and mounted with mounting medium (Permount^™^, Cat. No. SP15-100, Fisher Chemical) for better preservation.

### Glycosaminoglycan (GAG) and DNA quantifications

Sulfated glycosaminoglycans (sulfated GAGs) were quantified using the GAG Assay Kit (Cat. No. 6022, Chondrex) according to the manufacturer’s protocol. Briefly, an isolated trachea sample was weighed and digested in papain solution (concentration: 125 μg.mL^-1^; Cat. No. 60224, Chondrex^®^) at 60°C overnight. The absorbance of the solution was measured at 525 nm using a microplate reader (Synergy HTX multi-mode reader, BioTekTM), and the measured values were normalized to the wet weight of the tissue sample. Similarly, DNA contents were measured using the Quant-it PicoGreen double-stranded DNA Kit (Quant-iT^™^ PicoGreen^™^ dsDNA assay kit, Cat. No. P7589, Invitrogen). Briefly, tissue isolate was digested in papain solution (concentration: 125 μg.mL^-1^) for 4 hours at 60°C and the fluorescence intensity of the solution was measured (excitation/emission: 480 nm/530 nm) using the microplate reader. A standard calibration curve was produced by bacteriophage lambda DNA to quantify the DNA content in the solution.

### Scanning electron microscopy (SEM)

The luminal surfaces of the tracheas were visualized via scanning electron microscopy (SEM). Both native (control) and de-epithelialized tracheas were cut into small tissue samples (length: ~5 mm), which were then fixed in 2.5% glutaraldehyde at pH 7.4 at 4°C overnight. The fixed tissues were washed three times in 1× PBS and dehydrated in graded series of aqueous ethanol solutions (concentrations: 35, 50, 70, 85, 95, and 100% v/v). The tissues were kept in hexamethyldisilazane solvent (HMDS, Cat. No. AC120585000, Fisher Scientific) and air-dried overnight. The dried tissues were coated with gold using a sputter coater (EM MED020, Leica) which were then imaged using a scanning electron microscope (SEM; Auriga 40, Zeiss) with an accelerating voltage of 2 kV.

### Chondrocyte viability

To assess the viability of chondrocytes after de-epithelization and long-term culture, native (control) de-epithelized trachea (treated with 4% SDS solution) was cultured for two weeks in the bioreactor. After de-epithelialization, the luminal and exterior surfaces of the trachea were rinsed with the complete culture medium. The bioreactor was then transferred into an incubator at 37°C and 5% CO2. The culture medium in the trachea lumen and within the chamber was refreshed via a syringe pump (flow rate: 5 mL.h^-1^; volume: 10 mL; AL-4000, World Precision Instruments) and a programmable pump (flow rate: 5 mL.min-1; L/S^®^ standard digital pump system, Cole-Parmer), respectively, every day. After two weeks, the trachea was sectioned and stained with live/dead reagent staining (LIVE/DEAD^™^ Viability/Toxicity Kit, Cat. No. L3224 Invitrogen) according to manufacturer instruction. Briefly, the trachea samples were cut into slices (thickness ~500 μm) and then incubated in 1 mL 1× PBS containing 10 μM calcein AM and 20 μM ethidium homodimer-1 (EthD-1) for 15 min. After incubation, 1× PBS was used to rinse the samples. A fluorescent microscope (D-Eclipse C1, Nikon) equipped with fluorescent optical filters with excitation/emission of 495nm/515nm for live cells and excitation/emission of 495nm/635nm for dead cells was used to assess the viability of cartilage in de-epithelialized tracheas.

### Cytotoxicity evaluation of the de-epithelialized trachea

To evaluate the cytotoxicity of the de-epithelialized tracheas, mesenchymal stem cells (MSCs) were seeded and cultured on the luminal surface of the tracheas in the bioreactor. Survival and growth of the seeded cells were assessed during one week of *in vitro* culture. To do this, the de-epithelialized rat trachea treated with 4% SDS solution in the bioreactor was rinsed thoroughly using the culture medium. Next, the CFSE-labeled MSCs were suspended in the cell culture medium (cell concentration: 50,000 cells.mL^-1^; volume: 100 μL) were introduced into the trachea to deposit the cells onto the luminal surface. The trachea-loaded bioreactor was then placed in a gas-controlled cell incubator (MCO-20AIC, Panasonic) to culture the cell-seeded trachea at 37°C and 5% CO2 for 1, 4, or 7 days. During the cell culture, the culture medium within the inner space of the trachea was replaced with the fresh medium once a day using a programmable syringe pump (flow rate: 5 mL.h^-1^; volume: 10 mL; AL-4000, World Precision Instruments) and the removed culture medium was collected in a waste reservoir. In the meantime, the culture medium within the tissue culture chamber of the bioreactor was replaced continuously by circulating the medium between a reservoir and the chamber using a peristaltic pump (flow rate: 5 mL.min^-1^; L/S^®^ standard digital pump system, Cole-Parmer). The culture medium in the reservoir (total volume: 50 mL) was replaced once in two days. Viability and proliferation of the trachea seeded with CFSE-labelled MSCs were evaluated following 1, 4, or 7 days of *in vitro* culture by measuring fluorescence signal emitted from the cells (excitation/emission: 492 nm/518 nm) using a fluorescent microscope (Eclipse E1000, Nikon) or our micro-imaging device.

### Study design and statistical analysis

We present all data as mean ± standard deviation (SD). All experiments were conducted at least three times to **ensure** reproducibility of the results and to evaluate statistical significance within each experimental group. The de-epithelialized tracheas were compared with healthy native trachea freshly isolated from Sprague-Dawley rats. One-way analysis of variance (One-way ANOVA) was used following by Tukey’s Honest Significant Post-hoc test to compare the means of control groups. We considered a difference statistically significant if *p* < 0.05.

## Author Contributions

S.M.M, J.C., and J.K. conceived the concept of this work. S.M.M, J.C., and J.K. designed the experiments. S.M.M and J.C. performed the experiments. S.M.M, J.C., M.R.P., J.D.O., S.X.L.H., G.V.N., and J.K. analyzed the results. S.M.M., M.R.P., J.D.O., G.V.N., and J.K. wrote the manuscript.

## Acknowledgements

This research has been supported in part by American Thoracic Society Foundation Research Program and Research Grants from New Jersey Health Foundation provided to J.K. and the National Institutes of Health (P41 EB027062) to G.V.N. The authors would like to thank Drs. Tsengming (Alex) Chou and Matthew Libera, and Mr. Xinpei Wu at Stevens Laboratory for Multiscale Imaging (LMSI) for their assistance with SEM imaging of the tissue samples.

## References

1. F. Li, J. He, J. Wei, W. C. Cho and X. Liu, Stem Cells Int., 2015, 2015, 728307.

2. R. E. Rayner, P. Makena, G. L. Prasad and E. Cormet-Boyaka, Sci. Rep., 2019, 9, 500.

3. C. H. Dean and R. J. Snelgrove, Am. J. Respir. Crit. Care Med., 2018, 198, 1355–1356.

4. R. G. Crystal, S. H. Randell, J. F. Engelhardt, J. Voynow and M. E. Sunday, Proc. Am. Thorac. Soc., 2008, 5, 772–777.

5. A. Wanner, M. Salathé and T. G. O’Riordan, Am. J. Respir. Crit. Care Med., 1996, 154, 1868–1902.

6. C. Coraux, R. Hajj, P. Lesimple and E. Puchelle, Eur. Respir. Rev., 2005, 14, 131.

7. A. R. Berkebile, J. A. Bartlett, M. Abou Alaiwa, S. M. Varga, U. F. Power and P. B. McCray, Am. J. Respir. Cell Mol. Biol., 2019, 62, 104–111.

8. M. Cohen-Cymberknoh, E. Kerem, T. Ferkol and A. Elizur, Thorax, 2013, 68, 1157.

9. J. Mianné, E. Ahmed, C. Bourguignon, M. Fieldes, I. Vachier, A. Bourdin, S. Assou and J. De Vos, Am. J. Respir. Cell Mol. Biol., 2018, 59, 672–683.

10. E. Roscioli, R. Hamon, S. E. Lester, H. P. A. Jersmann, P. N. Reynolds and S. Hodge, Respir. Res., 2018, 19, 234.

11. N. T. N. Trinh, O. Bardou, A. Privé, E. Maillé, D. Adam, S. Lingée, P. Ferraro, M.-Y. Desrosiers, C. Coraux and E. Brochiero, Eur. Respir. J., 2012, 40, 1390.

12. A. Horani and T. W. Ferkol, Chest, 2018, 154, 645–652.

13. C. H. Hochberg and V. K. Sidhaye, in Lung Epithelial Biology in the Pathogenesis of Pulmonary Disease, eds. V. K. Sidhaye and M. Koval, Academic Press, Boston, 2017, pp. 165–184.

14. K. H. Benam, R. Villenave, C. Lucchesi, A. Varone, C. Hubeau, H.-H. Lee, S. E. Alves, M. Salmon, T. C. Ferrante, J. C. Weaver, A. Bahinski, G. A. Hamilton and D. E. Ingber, Nat. Methods, 2016, 13, 151–157.

15. S. E. Gilpin, J. M. Charest, X. Ren and H. C. Ott, Thorac. Surg. Clin., 2016, 26, 163–171.

16. M. Z. Nikolić, O. Caritg, Q. Jeng, J. A. Johnson, D. Sun, K. J. Howell, J. L. Brady, U. Laresgoiti, G. Allen, R. Butler, M. Zilbauer, A. Giangreco and E. L. Rawlins, Elife, 2017, 6.

17. B. F. Niemeyer, P. Zhao, R. M. Tuder and K. H. Benam, SLAS Discov., 2018, 23, 777–789.

18. K. B. McCauley, F. Hawkins, M. Serra, D. C. Thomas, A. Jacob and D. N. Kotton, Cell Stem Cell, 2017, 20, 844–857.e846.

19. R. Plebani, R. Potla, M. Soong, H. Bai, Z. Izadifar, A. Jiang, R. N. Travis, C. Belgur, M. J. Cartwright, R. Prantil-Baun, P. Jolly, S. E. Giplin, M. Romano and D. E. Ingber, medRxiv, 2021, DOI: 10.1101/2021.07.15.21260407, 2021.2007.2015.21260407.

20. L. E. Haswell, K. Hewitt, D. Thorne, A. Richter and M. D. Gaça, Toxicol. In Vitro, 2010, 24, 981–987.

21. C. Xiao, S. M. Puddicombe, S. Field, J. Haywood, V. Broughton-Head, I. Puxeddu, H. M. Haitchi, E. Vernon-Wilson, D. Sammut, N. Bedke, C. Cremin, J. Sones, R. Djukanović, P. H. Howarth, J. E. Collins, S. T. Holgate, P. Monk and D. E. Davies, J. Allergy Clin. Immunol., 2011, 128, 549–556.e512.

22. A. Jacob, M. Morley, F. Hawkins, K. B. McCauley, J. C. Jean, H. Heins, C. L. Na, T. E. Weaver, M. Vedaie, K. Hurley, A. Hinds, S. J. Russo, S. Kook, W. Zacharias, M. Ochs, K. Traber, L. J. Quinton, A. Crane, B. R. Davis, F. V. White, J. Wambach, J. A. Whitsett, F. S. Cole, E. E. Morrisey, S. H. Guttentag, M. F. Beers and D. N. Kotton, Cell Stem Cell, 2017, 21, 472–488.e410.

23. A. Strikoudis, A. Cieślak, L. Loffredo, Y.-W. Chen, N. Patel, A. Saqi, D. J. Lederer and H.-W. Snoeck, Cell Rep., 2019, 27, 3709–3723.e3705.

24. E. Schruf, V. Schroeder, H. Q. Le, T. Schönberger, D. Raedel, E. L. Stewart, K. Fundel-Clemens, T. Bluhmki, S. Weigle, M. Schuler, M. J. Thomas, R. Heilker, M. J. Webster, M. Dass, M. Frick, B. Stierstorfer, K. Quast and J. P. Garnett, The FASEB Journal, 2020, 34, 7825–7846.

25. H. Surendran, S. Nandakumar and R. Pal, Stem Cells Dev., 2020, 29, 1365–1369.

26. S. X. Huang, M. N. Islam, J. O’Neill, Z. Hu, Y. G. Yang, Y. W. Chen, M. Mumau, M. D. Green, G. Vunjak-Novakovic, J. Bhattacharya and H. W. Snoeck, Nat. Biotechnol., 2014, 32, 84–91.

27. A. J. Miller, B. R. Dye, D. Ferrer-Torres, D. R. Hill, A. W. Overeem, L. D. Shea and J. R. Spence, Nat. Protoc., 2019, 14, 518–540.

28. F. J. Hawkins, S. Suzuki, M. L. Beermann, C. Barillà, R. Wang, C. Villacorta-Martin, A. Berical, J. C. Jean, J. Le Suer, T. Matte, C. Simone-Roach, Y. Tang, T. M. Schlaeger, A. M. Crane, N. Matthias, S. X. L. Huang, S. H. Randell, J. Wu, J. R. Spence, G. Carraro, B. R. Stripp, A. Rab, E. J. Sorsher, A. Horani, S. L. Brody, B. R. Davis and D. N. Kotton, Cell Stem Cell, 2021, 28, 79–95.e78.

29. P. Duchesneau, T. K. Waddell and G. Karoubi, Int. J. Mol. Sci., 2020, 21, 5219.

30. B. A. Guenthart, J. D. O’Neill, J. Kim, K. Fung, G. Vunjak-Novakovic and M. Bacchetta, J. Heart Lung Transplant., 2019, 38, 215–224.

31. L. Van Haute, G. De Block, I. Liebaers, K. Sermon and M. De Rycke, Respir. Res., 2009, 10, 105.

32. S. Halldorsson, V. Asgrimsson, I. Axelsson, G. H. Gudmundsson, M. Steinarsdottir, O. Baldursson and T. Gudjonsson, In Vitro Cell. Dev. Biol. Anim., 2007, 43, 283–289.

33. H. Lin, H. Li, H.-J. Cho, S. Bian, H.-J. Roh, M.-K. Lee, J. S. Kim, S.-J. Chung, C.-K. Shim and D.-D. Kim, J. Pharm. Sci., 2007, 96, 341–350.

34. J. C. Nawroth, R. Barrile, D. Conegliano, S. van Riet, P. S. Hiemstra and R. Villenave, Adv Drug Deliv Rev, 2019, 140, 12–32.

35. K. Gkatzis, S. Taghizadeh, D. Huh, D. Y. R. Stainier and S. Bellusci, Eur. Respir. J., 2018, 52.

36. D. Huh, H. J. Kim, J. P. Fraser, D. E. Shea, M. Khan, A. Bahinski, G. A. Hamilton and D. E. Ingber, Nat. Protoc., 2013, 8, 2135–2157.

37. L. Si, H. Bai, M. Rodas, W. Cao, C. Y. Oh, A. Jiang, R. Moller, D. Hoagland, K. Oishi, S. Horiuchi, S. Uhl, D. Blanco-Melo, R. A. Albrecht, W.-C. Liu, T. Jordan, B. E. Nilsson-Payant, I. Golynker, J. Frere, J. Logue, R. Haupt, M. McGrath, S. Weston, T. Zhang, R. Plebani, M. Soong, A. Nurani, S. M. Kim, D. Y. Zhu, K. H. Benam, G. Goyal, S. E. Gilpin, R. Prantil-Baun, S. P. Gygi, R. K. Powers, K. E. Carlson, M. Frieman, B. R. tenOever and D. E. Ingber, Nat. Biomed. Eng., 2021, DOI: 10.1038/s41551-021-00718-9.

38. K. H. Benam, M. Mazur, Y. Choe, T. C. Ferrante, R. Novak and D. E. Ingber, Methods Mol. Biol., 2017, 1612, 345–365.

39. K. Hajipouran Benam, R. Novak, R. Villenave, C. Lucchesi, C. Hubeau, J. Nawroth, M. Hirano-Kobayashi, T. C. Ferrante, Y. Choe, R. Prantil-Baun, J. C. Weaver, G. A. Hamilton, A. Bahinski and D. E. Ingber, Eur. Respir. J., 2016, 48, OA4542.

40. L. G. Griffith and M. A. Swartz, Nature Reviews Molecular Cell Biology, 2006, 7, 211–224.

41. A. M. Greaney, T. S. Adams, M. S. Brickman Raredon, E. Gubbins, J. C. Schupp, A. J. Engler, M. Ghaedi, Y. Yuan, N. Kaminski and L. E. Niklason, Cell Rep., 2020, 30, 4250–4265.e4256.

42. M. D. Green, S. X. Huang and H. W. Snoeck, Bioessays, 2013, 35, 261–270.

43. C. R. Butler, R. E. Hynds, K. H. Gowers, H. Lee Ddo, J. M. Brown, C. Crowley, V. H. Teixeira, C. M. Smith, L. Urbani, N. J. Hamilton, R. M. Thakrar, H. L. Booth, M. A. Birchall, P. De Coppi, A. Giangreco, C. O’Callaghan and S. M. Janes, Am J Respir Crit Care Med, 2016, 194, 156–168.

44. M. J. Goldman, Y. Yang and J. M. Wilson, Nat. Genet., 1995, 9, 126–131.

45. D. W. Borthwick, M. Shahbazian, Q. T. Krantz, J. R. Dorin and S. H. Randell, Am J Respir Cell Mol Biol, 2001, 24, 662–670.

46. R. Hajj, T. Baranek, R. Le Naour, P. Lesimple, E. Puchelle and C. Coraux, Stem Cells, 2007, 25, 139–148.

47. S. X. L. Huang, M. D. Green, A. T. de Carvalho, M. Mumau, Y.-W. Chen, S. L. D’Souza and H.-W. Snoeck, Nat. Protoc., 2015, 10, 413–425.

48. R. Nossa, J. Costa, L. Cacopardo and A. Ahluwalia, Journal of Tissue Engineering, 2021, 12, 20417314211008696.

49. J. Kim, J. D. O’Neill, N. V. Dorrello, M. Bacchetta and G. Vunjak-Novakovic, Proc. Natl. Acad. Sci. U. S. A., 2015, 112, 11530–11535.

50. J. Kim, B. Guenthart, J. D. O’Neill, N. V. Dorrello, M. Bacchetta and G. Vunjak-Novakovic, Sci. Rep., 2017, 7, 13082.

51. J. Kim, J. D. O’Neill and G. Vunjak-Novakovic, Appl. Phys. Lett., 2015, 107, 144101.

52. J. L. Balestrini, A. L. Gard, A. Liu, K. L. Leiby, J. Schwan, B. Kunkemoeller, E. A. Calle, A. Sivarapatna, T. Lin, S. Dimitrievska, S. G. Cambpell and L. E. Niklason, Integr. Biol., 2015, 7, 1598–1610.

53. D. A. Taylor, L. C. Sampaio, Z. Ferdous, A. S. Gobin and L. J. Taite, Acta Biomater., 2018, 74, 74–89.

54. J. L. Balestrini, A. L. Gard, A. Liu, K. L. Leiby, J. Schwan, B. Kunkemoeller, E. A. Calle, A. Sivarapatna, T. Lin, S. Dimitrievska, S. G. Cambpell and L. E. Niklason, Integrative biology: quantitative biosciences from nano to macro, 2015, 7, 1598–1610.

55. L. M. Godin, B. J. Sandri, D. E. Wagner, C. M. Meyer, A. P. Price, I. Akinnola, D. J. Weiss and A. Panoskaltsis-Mortari, PLoS One, 2016, 11, e0150966.

56. N. Qu, P. de Vos, M. Schelfhorst, A. de Haan, W. Timens and J. Prop, J. Heart Lung Transplant., 2005, 24, 882–890.

57. M. Zang, Q. Zhang, E. I. Chang, A. B. Mathur and P. Yu, Plast. Reconstr. Surg., 2013, 132.

58. Z. Dimou, E. Michalopoulos, M. Katsimpoulas, D. Dimitroulis, G. Kouraklis, C. Stavropoulos-Giokas and P. Tomos, Anticancer Res., 2019, 39, 145–150.

59. S. Haykal, M. Salna, Y. Zhou, P. Marcus, M. Fatehi, G. Frost, T. Machuca, S. O. Hofer and T. K. Waddell, Tissue Eng Part C Methods, 2014, 20, 681–692.

60. P. Hong, M. Bezuhly, M. E. Graham and P. F. Gratzer, Int. J. Pediatr. Otorhinolaryngol., 2018, 112, 67–74.

