## Supplementary Information for "Imaging-Guided Bioreactor for De-Epithelialization and Long-Term Cultivation of *Ex Vivo* Rat Trachea"

### I. Supplementary Methods and Materials

#### 1. Image processing of the mechanical vibration videos

High-frame rate imaging method was used to monitor the movement of the mechanical vibration stage. To process the videos, we used the methods described in our previous report [S1]. Briefly, videos recorded at 240 frame per second (fps) were stored in AVI file format which is a standard video file format compatible with ImageJ. To determine the displacement distance of the stage, a ruler was placed adjacent to the shaker as a reference object with known length. The video file was imported to ImageJ as a sequence of image frames using "AVI reader" plugin. The displacement distance of the sample stage in the image sequence was measured with respect to the reference ruler. In the meantime, to improve visibility of the stage movement, "enhance contrast" and "find edges" functions were used in ImageJ. To plot the displacement curves, a small region of the stage moving up and down was cropped and extracted from each image frame. The cropped images were then stitched horizontally with "grid/collection stitching" plugin as a single image file. The similar procedure was done for different regions of the stage to generate a continuous waveform that represent displacement of the sample stage over time.

### II. Supplementary Figures

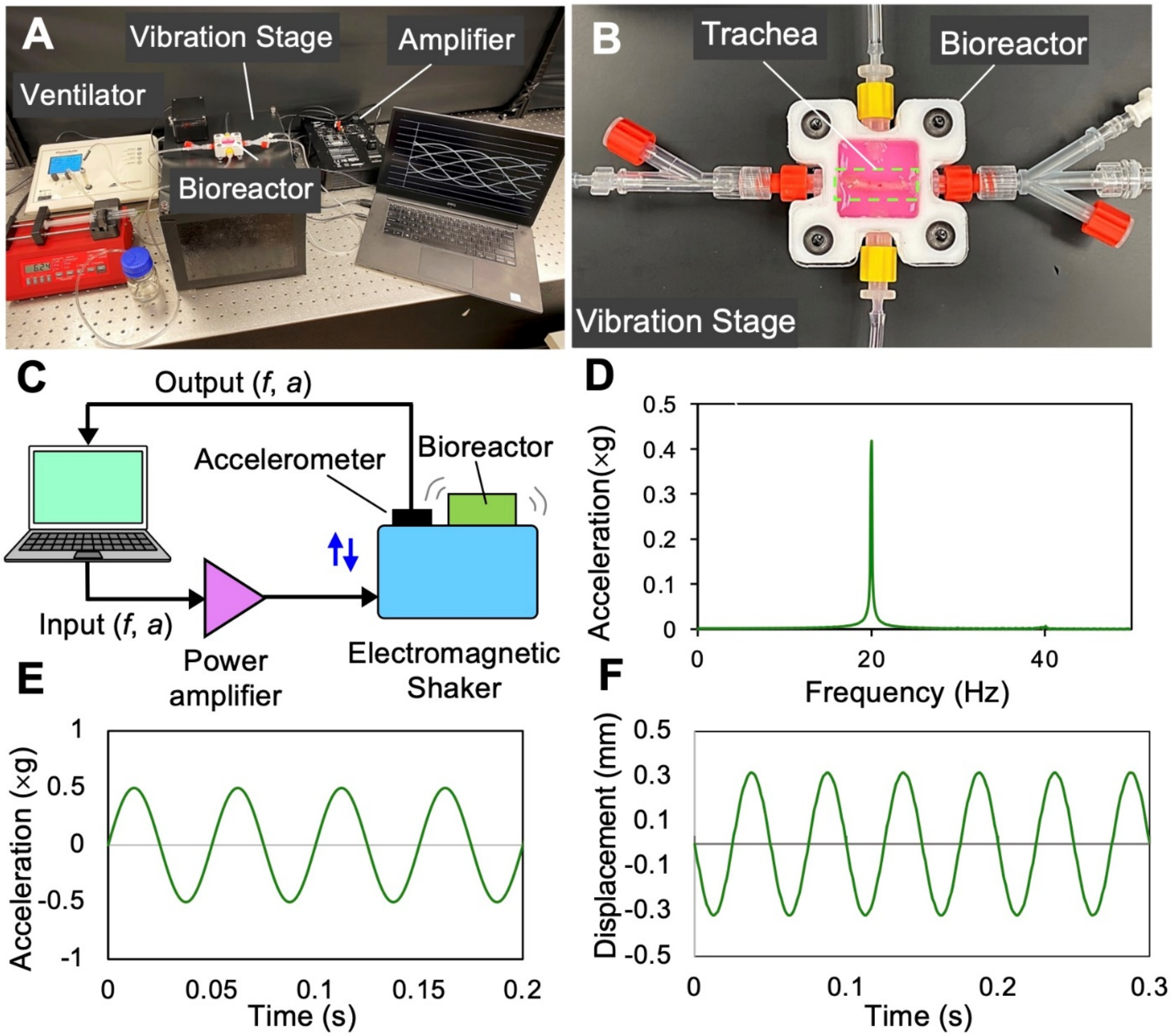


**Figure S1. Mechanical vibration for clearance of detergent-disrupted epithelial cells:** (**A**) After cells lysis by SDS detergent solution, the bioreactor containing the trachea was mechanically vibrated using a custom-built electromagnetic shaker. (**B**) Photograph showing the trachea bioreactor placed on the shaker. (**C**) Schematic of the electromagnetic shaker where the oscillation pattern (e.g., *f*: frequency, *a*: amplitude) was controlled by a computerized waveform generator and response was monitored via an accelerometer and a high-frame rate camera. (**D**) Acceleration force measured against frequency showed the maximum force generation at 20 Hz which was the input frequency of the waveform used for tissue vibration. (**E**) Output acceleration force measured against time and (**F**) displacement of the sample stage of the shaker determined using a high-speed camera and subsequent image processing via ImageJ (**Video S1**).

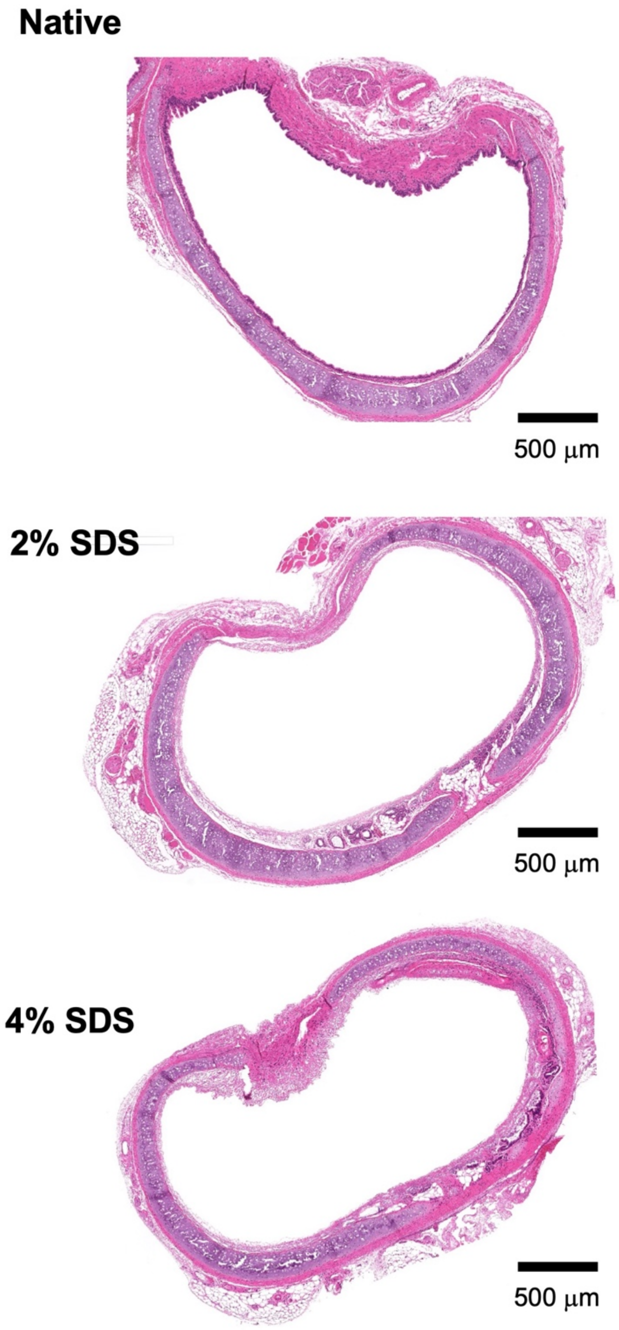


**Figure S2.** H&E staining images of the cross-section of native and de-epithelialized rat tracheas. De-epithelialization was achieved by topical deposition of 2% and 4% SDS detergent solution followed by vibration-assisted airway wash.

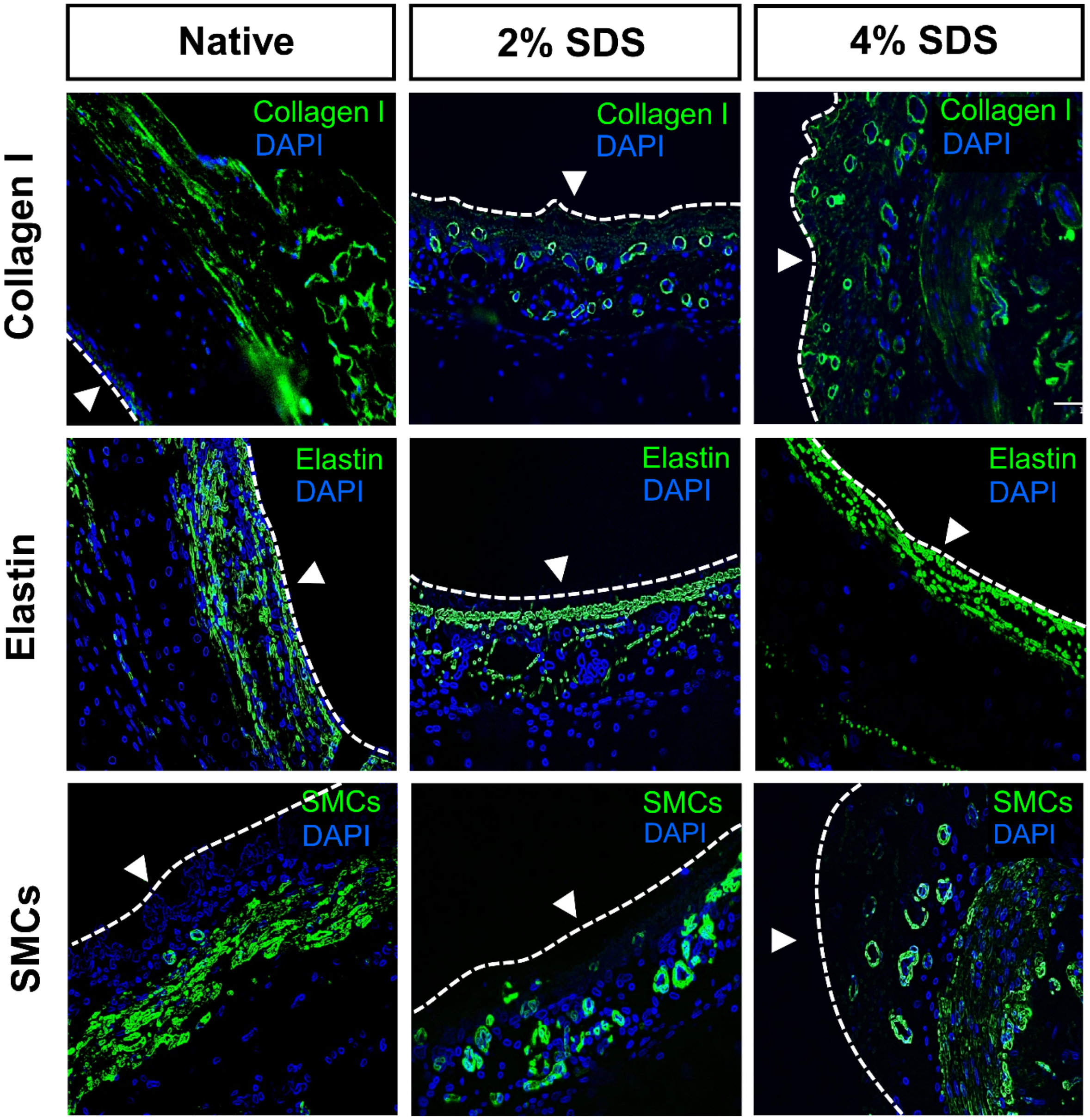


**Figure S3.** Immunofluorescence staining of the rat trachea tissues with collagen I, elastin, and smooth muscles (SMCs). Discernible green signals were observed in the tissues that were de-epithelialized with SDS detergent solutions, indicating preservation of the ECM components and structures. Tracheal lumen is indicated by an arrowhead in each image.

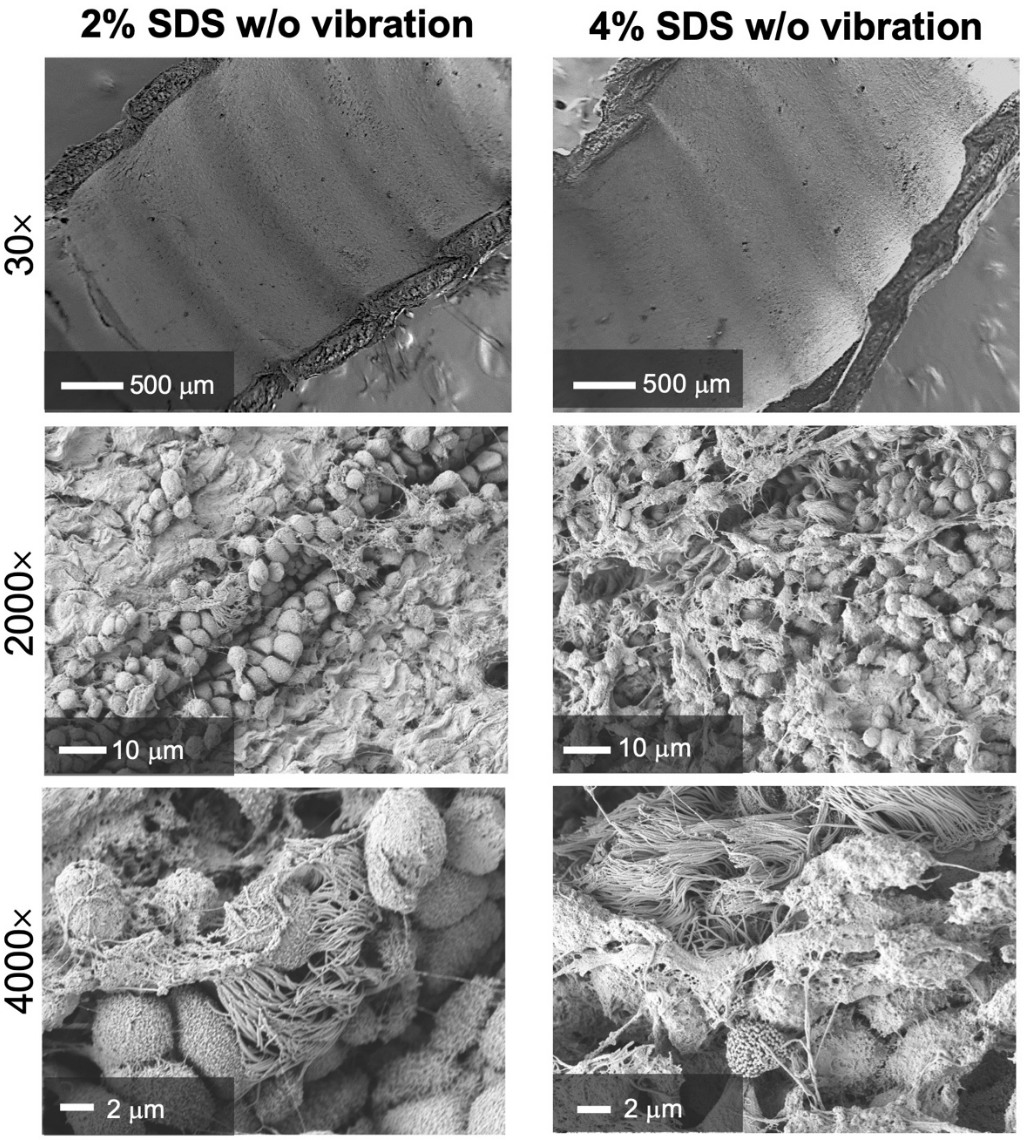


**Figure S4.** SEM images showing rat tracheas treated with 2% and 4% SDS solution followed by gentle airway washing in the absence of mechanical vibration. Debris of epithelial cells remained adhered onto the lumen of these tracheas suggested that vibration energy provided to the tissue during de-epithelialization was essential to achieve complete epithelium removal from the airway lumen.

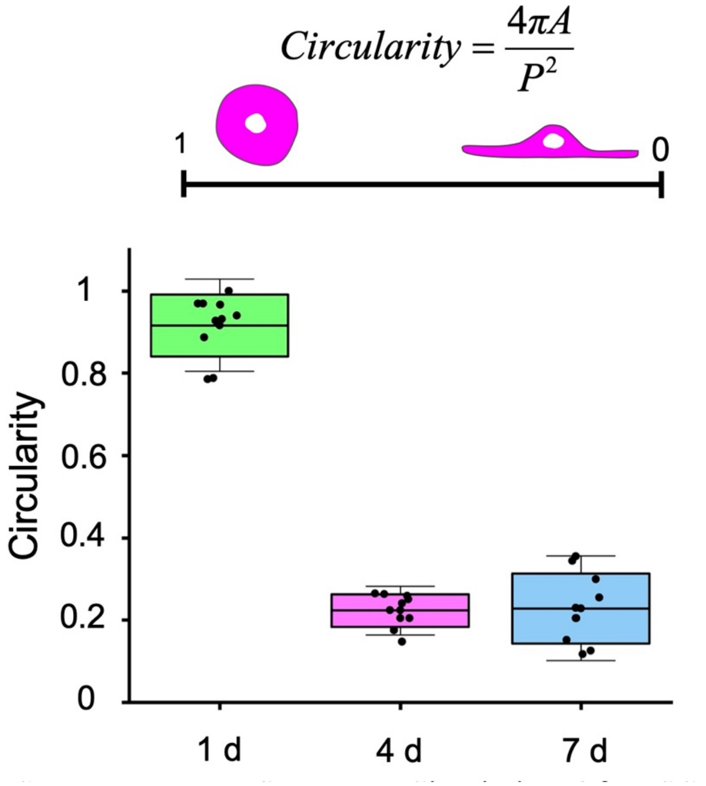


**Figure S5.** Circularity of MSCs implanted and cultured on the de-epithelialized tracheas measured at different time intervals (i.e., 1, 4, and 7 days). Cell circularity (range: 0-1) was determined by calculating the ratio of the surface area (*A*) to the perimeter (*P*) of the cell. Circularity of the cells decreased substantially to below 0.25 at days 4 and 7, indicating the cells were actively engrafted onto the tissue surface.

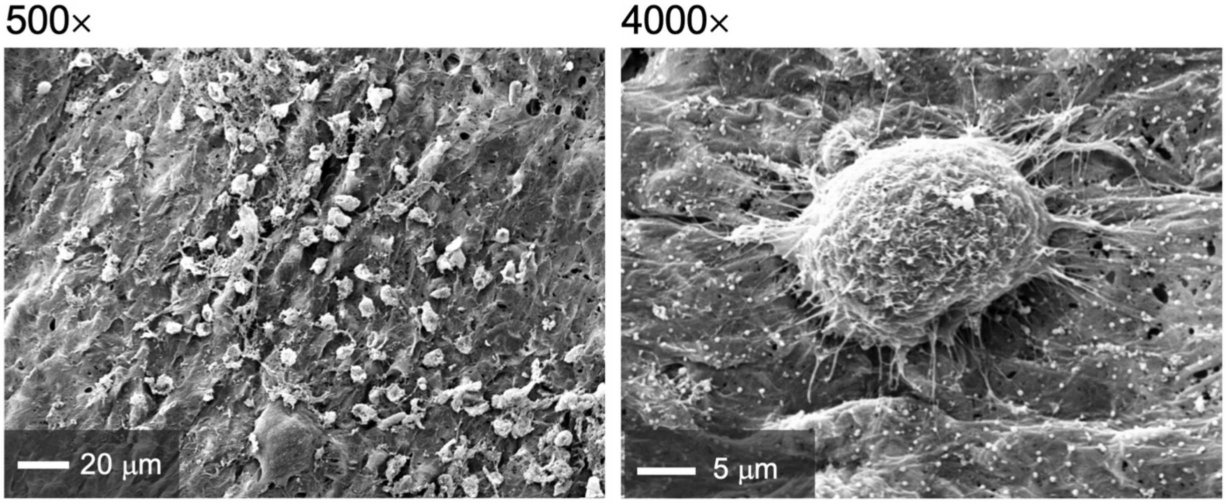


**Figure S6.** SEM micrographs of MSCs distributed onto the de-epithelialized rat trachea lumen. Notably, the cells initiated binding onto the tissue surface via multiple protrusions extending from the cell membrane to the surface at day 1.

### III. Supplementary Tables

**Table S1.** List of primary antibodies.

| Primary antibodies | Application | Catalog number, Company | Dilution |
|---|---|---|---|
| Rb Anti-Elastin | IF | ab21610, Abcam | 1:250 |
| Rb Anti-EpCAM | IF | PA5-19832, ThermoFisher | 1:1000 |
| Rb Anti-Collagen I | IF | ab34710, Abcam | 1:200 |
| Rb Anti-Alpha Smooth Muscle actin | IF | ab124964, Abcam | 1:200 |
| Rb anti-Laminin | IF | PA1-16730, ThermoFisher | 1:1000 |
| Rb Anti-CD31 | IF | ab28364, Abcam | 1:200 |
| Mouse Anti-Rat CD31 | IF | MCA1334GA, BioRad | 1:100 |

*IF: immunofluorescence

**Table S2.** List of secondary antibodies.

| Secondary antibodies | Application | Catalog number, Company | Dilution |
|---|---|---|---|
| Donkey Anti-Rabbit IgG (488) | IF | ab150073, Abcam | 1:500 |
| Goat anti Mouse IgG (680) | IF | STAR117D680GA, BioRad | 1:100 |
| Goat anti-Rabbit IgG (488) | IF | ab97244, Abcam | 1:300 |
| Goat anti-Rabbit IgG (555) | IF | ab150078, Abcam | 1:500 |

*IF: immunofluorescence

### IV. Legends for Supplementary Videos

**Video S1.** Movies showing the trachea-loaded bioreactor being mechanically vibrated on our custom-built shaker at 20 Hz of oscillation frequency. The videos were taken at 240 frame per second (fps) and being played at 30 fps.

**Video S2.** Side view of the sample stage of the shaker being oscillated at 20 Hz. The maximum vertical displacement was ± 0.3 mm with respect to the original position of the stage at still.

**Video S3.** Internal space of the isolated rat trachea visualized using the GRIN lens imaging probe while illuminating with white light.

**Video S4.** Internal space of the isolated rat trachea visualized using the GRIN lens imaging probe while illuminating with 488-nm laser. The tracheal lumen was stained with CFSE prior to the imaging and emission light was filtered using a bandpass filter (ET535/50 nm, Chroma®).
